# Effective utilization of xylose-rich substrates for butyric acid production by *Clostridium tyrobutyricum* ATCC 25755 despite carbon catabolite repression

**DOI:** 10.64898/2026.11.11.765203

**Authors:** Ki-Yeon Kim, Hyun Ju Oh, Jung Ho Ahn, Ja Kyong Ko, Gyeongtaek Gong, Youngsoon Um

## Abstract

Carbon catabolite repression (CCR) can restrict xylose utilization during butyric acid production from lignocellulosic hydrolysates by *Clostridium tyrobutyricum* ATCC 25755. This study evaluated CCR behavior using 2-deoxy-D-glucose (2-DG), a glucose analog, and examined the effects of glucose–xylose composition on mixed-sugar fermentation. 2-DG caused concentration-dependent growth inhibition in xylose-containing medium, indicating CCR-like behavior. However, xylose was partially co-consumed in the presence of glucose or 2-DG, suggesting incomplete inhibition. In batch fermentations with 80 g/L total sugars, the overall xylose consumption rate declined from 2.68 to 0.53 g/L/h as the initial glucose concentration increased from 20 to 60 g/L. Nevertheless, a xylose-rich mixture (20 g/L glucose and 60 g/L xylose) achieved an overall total sugar consumption rate of 3.54 g/L/h and a butyric acid productivity of 1.22 g/L/h, comparable to those with 80 g/L glucose alone (3.64 and 1.26 g/L/h, respectively). Detectable xylose isomerase activity during glucose– xylose co-utilization further supported incomplete repression of xylose metabolism. In xylose-rich rice straw hydrolysate (5.0 g/L glucose and 15.1 g/L xylose), both sugars were completely consumed, producing 7.68 g/L butyric acid. These findings suggest that xylose-rich mixtures can support effective sugar utilization and butyric acid production by *C. tyrobutyricum* despite its CCR-like behavior.

## 1. Introduction

Concerns over climate change associated with fossil-derived CO₂ emissions have increased interest in the biological production of fuels and chemicals from abundant, renewable lignocellulosic biomass (Saravanan et al., 2022). Pretreatment and hydrolysis of lignocellulosic biomass release fermentable hexose and pentose sugars, of which glucose and xylose are major components (Suo et al., 2018). Because many microorganisms preferentially utilize glucose, effective xylose utilization is important for improving the conversion of lignocellulosic sugars into desired products. *Clostridium tyrobutyricum* ATCC 25755 is a promising candidate for butyric acid production because it can metabolize either glucose or xylose as the sole carbon source with high product selectivity and yield (Bao et al., 2020; Fu et al., 2017; Jiang et al., 2010; Zhu et al., 2002).

Butyric acid production from glucose–xylose mixtures and lignocellulosic hydrolysates has been reported using wild-type *C. tyrobutyricum* ATCC 25755 (Fu et al., 2017; Fu et al., 2022; Oh et al., 2019; Suo et al., 2018). However, glucose-mediated inhibition of xylose utilization can limit sugar conversion in mixed-substrate fermentations. This inhibition is associated with carbon catabolite repression (CCR), a regulatory phenomenon through which a preferred carbon source suppresses the utilization of alternative carbon sources (Görke and Stülke, 2008). Several metabolic engineering approaches have therefore been investigated to alleviate CCR in *C. tyrobutyricum* ATCC 25755. Fu et al. (2017) overexpressed three heterologous genes involved in xylose transport and metabolism, *xylT*, *xylA*, and *xylB*, from *C. acetobutylicum* to enable simultaneous glucose and xylose utilization. Fu et al. (2022) subsequently demonstrated contributions from both CcpA-mediated regulation and XylR-mediated repression and reported that combined inactivation of *hprK* and *xylR* eliminated glucose-mediated repression under the conditions examined.

Although *C. tyrobutyricum* ATCC 25755 exhibits CCR through the regulatory mechanisms described above, the extent of glucose-mediated repression appears to vary with fermentation conditions. Pronounced preferential glucose consumption and limited xylose consumption while glucose remained available have been reported in wild-type cultures supplied with synthetic glucose–xylose mixtures (Fu et al., 2017; Fu et al., 2022; Suo et al., 2018). In contrast, partial glucose–xylose co-utilization has also been observed in fermentations using immobilized cells supplied with mixed sugars or lignocellulosic hydrolysates (Wei et al., 2013; Zhu et al., 2002), as well as in mixed-sugar fermentation using free cells (Oh et al., 2019). These differences in reported sugar consumption patterns highlight the need for further characterization of the extent of glucose-mediated repression in this strain.

The glucose analog 2-deoxy-D-glucose (2-DG) has been used to investigate CCR-related phenotypes and select strains with alleviated catabolite repression (Li & He, 2016; Liang et al., 2022; Tsakraklides et al., 2012). Unlike glucose, 2-DG lacks a hydroxyl group at the second carbon position and, following cellular uptake and phosphorylation, is generally not efficiently metabolized through glycolysis. The addition of 2-DG can inhibit growth on non-preferred carbon sources such as xylose and has therefore been used to examine CCR-like responses (Liang et al., 2022; Tsakraklides et al., 2012). Growth on xylose in the presence of 2-DG has also been used to select variants with improved mixed-sugar utilization (Li & He, 2016; Liang et al., 2022). Thus, 2-DG provides a complementary approach to examining inhibition by a glucose analog without supplying glucose as a metabolizable substrate.

To assess the implications of CCR for mixed-sugar fermentation, sugar utilization must also be examined when glucose and xylose are supplied together. Glucose-to-xylose ratios in lignocellulosic hydrolysates vary with the feedstock and pretreatment and hydrolysis conditions (Fu et al., 2022; Liang et al., 2013). Examining how these ratios influence xylose consumption and butyric acid production may therefore help determine the extent to which glucose-mediated inhibition limits fermentation performance. Accordingly, evaluating 2-DG responses together with sugar consumption profiles at different glucose-to-xylose ratios can provide a more informative assessment of CCR-like behavior and its implications for mixed-sugar fermentation.

In this study, the effects of 2-DG on the growth of *C. tyrobutyricum* ATCC 25755 with xylose as the sole carbon source were examined to assess CCR behavior. Growth and sugar consumption were subsequently evaluated in glucose–xylose mixtures with and without 2-DG. The effects of different glucose-to-xylose ratios on sugar utilization and butyric acid production were investigated, and xylose isomerase activity was measured to assess whether xylose metabolism remained active in the presence of glucose. Fermentation of xylose-rich rice straw hydrolysate was then evaluated to examine the applicability of the observed consumption patterns to a lignocellulosic substrate. Together, these experiments aimed to clarify the fermentation behavior of this strain and identify conditions that support effective utilization of xylose-rich substrates without genetic modification to eliminate CCR.

## 2. Materials and methods

### 2.1. Bacterial strain, medium, and culture conditions

*C. tyrobutyricum* ATCC 25755 was obtained from the American Type Culture Collection (Manassas, VA, USA). Modified reinforced clostridial medium (RCM), containing 10 g/L tryptone, 10 g/L beef extract, 3 g/L yeast extract, 5 g/L glucose, 5 g/L NaCl, 1 g/L soluble starch, 0.5 g/L L-cysteine HCl, and 3.0 g/L sodium acetate, was used to prepare glycerol stocks and seed cultures. Glycerol stocks containing 25% glycerol were stored at –80 °C. Modified P2 (mP2) medium was used for the fermentation experiments. The mP2 medium contained 5.0 g/L yeast extract, 0.2 g/L MgSO_4_, 0.01 g/L MnSO_4_, 0.01 g/L FeSO_4_, 0.01 g/L NaCl, 0.5 g/L KH_2_PO_4_, 0.5 g/L K_2_HPO_4_, 2.0 g/L ammonium acetate, 0.001 g/L resazurin, and 100 mM MES (2-(*N*-morpholino)ethanesulfonic acid hydrate) for pH buffering. The initial pH of the medium was adjusted to 6.5 with 4 M KOH.

For serum bottle experiments, glucose, xylose, and 2-DG were added at the concentrations specified in Section 2.2. Serum bottles with a total volume of 157 mL were filled with 50 mL medium, purged with argon gas, and sealed with butyl rubber stoppers and aluminum crimp seals. After autoclaving at 121 °C for 20 min, the medium was inoculated with 2.5% (v/v) seed culture and incubated at 37 °C and 150 rpm.

### 2.2. Effect of 2-deoxy-D-glucose (2-DG) on growth and sugar consumption

The effects of 2-deoxy-D-glucose (2-DG) on the growth of *C. tyrobutyricum* ATCC 25755 were examined to assess CCR behavior. To determine whether 2-DG could support growth, cultures were prepared with 0.5–10 g/L 2-DG in the absence of glucose and xylose. To examine its effects on growth in the presence of a metabolizable sugar, 2-DG was added at 0, 0.1, 0.5, 1.5, 3, 6, or 10 g/L to mP2 medium containing either 15 g/L glucose or 15 g/L xylose.

Growth and sugar consumption in glucose–xylose mixtures were also examined with and without 0.5 g/L 2-DG. Low initial concentrations of glucose (0.3–0.8 g/L) and xylose (1.0– 1.4 g/L) were used. All cultures were incubated at 37 °C and 150 rpm without pH control.

### 2.3. Batch fermentation in a 2.5 L bioreactor with pH control

Batch fermentations with different glucose-xylose compositions were conducted in a 2.5 L bioreactor (Fermentec, Korea) equipped with temperature and pH controllers. Synthetic sugar media contained a nominal total initial sugar concentration of 80 g/L, with the following glucose and xylose concentrations: 80 and 0 g/L (G80), 0 and 80 g/L (X80), 20 and 60 g/L (G20X60), 40 and 40 g/L (G40X40), and 60 and 20 g/L (G60X20), respectively. Rice straw hydrolysate was also evaluated as a fermentation substrate.

The mP2 medium described in Section 2.1 was used, except that the yeast extract concentration was 25 g/L. The medium was purged with argon gas for 1 h before inoculation. The temperature was maintained at 37 °C, and the pH was automatically maintained at 6.0 by adding 6 N NaOH. The agitation speed was set to 200 rpm. Fermentations were initiated with 5% (v/v) seed culture at an OD₆₀₀ of 3–4, giving a final working volume of 1 L. Samples were collected periodically to monitor cell growth, sugar consumption, and butyric acid production.

### 2.4. Xylose isomerase activity

Samples were collected during the exponential and stationary phases, and cells were harvested by centrifugation (8,000 × g, 15 min, 4 °C). The pellets were washed twice with 50 mM sodium phosphate buffer (pH 6.0) and resuspended in the same buffer. Cells were disrupted by bead-beating, and the debris was removed by centrifugation (8,000 × g, 15 min, 4 °C). The supernatant was collected as the cell-free extract, and protein concentration was determined using the Bio-Rad Protein Assay Kit (Bio-Rad, USA).

Xylose isomerase activity was determined by measuring xylulose formation using the cysteine–sulfuric acid–carbazole method (Dische & Borenfreund, 1951). Briefly, 100 µL of cell-free extract was added to 500 µL of sodium phosphate buffer (pH 6.0) containing 1 mM MnCl₂ and 5 mM xylose, and incubated for 20 min at 37 °C. Formation of xylulose was quantified by measuring absorbance at 540 nm against a standard curve. Specific enzyme activity was expressed as U/mg protein, where one unit was defined as the production of 1 µmol xylulose per minute under the assay conditions.

### 2.5. Preparation of rice straw hydrolysate and fermentation

Dried rice straw was cut into ∼2 cm pieces and further milled using a laboratory disintegrator to obtain particles approximately 1 cm long and 1 mm thick. For delignification, the homogenized rice straw was treated with 0.2 N NaOH at a solid-to-liquid ratio of 1:10 (w/v) at room temperature for 4 h. The treated material was washed with tap water through a 0.3 mm sieve, and then dried at 55 °C to a constant weight. Acid hydrolysis was performed by immersing the dried matter in 1% H₂SO₄ at a solid-to-liquid ratio of 1:10 (w/v) and autoclaving at 121 °C for 80 min. After hydrolysis, solids were removed by filtration, and the pH of the filtrate was adjusted to 7.5 with Ca(OH)₂ while stirring. The mixture was incubated at room temperature for 1 h and filtered again. The rice straw hydrolysate was supplemented with the components of mP2 medium described in Section 2.1. The batch fermentation in the 2.5 L bioreactor was conducted under the conditions described in Section 2.3.

### 2.6. Analysis of cell growth, sugars, and butyric acid

The optical densities of cultures were measured at 600 nm with a UVmini-1240 Shimadzu spectrophotometer (Shimadzu, Tokyo, Japan). Sugar concentrations were determined by HPLC (Agilent 1260 series, Palo Alto, CA, USA) equipped with a refractive index detector (RID). Aliquots of culture collected at different time intervals were centrifuged at 10,000 × g for 10 min and the supernatants were transferred to new glass vials for analysis. For HPLC analysis of the supernatant, the Bio-Rad HPX-87-H column (30 cm × 7.8 mm) was used. The column temperature was maintained at 55 °C. The mobile phase was 0.05 N H_2_SO_4_, supplied at a flow rate of 0.5 mL/min. Butyric acid was analyzed using gas chromatography (Agilent Technologies, Model 6890, Palo Alto, CA, USA) equipped with a flame ionization detector (FID) and a 30 m × 0.25 mm × 0.25 µm HP-INNOWAX polyethylene glycol column.

## 3. Results and discussion

### 3.1. Effect of 2-DG on the growth of *C. tyrobutyricum* with xylose as the sole carbon source

2-DG, a glucose analog, has been used to investigate CCR because it can inhibit growth on alternative carbon sources such as xylose (Liang et al., 2022; Tsakraklides et al., 2012). Prior to examining whether 2-DG interfered with xylose utilization and consequently inhibited the growth of *C. tyrobutyricum* ATCC 25755, its ability to support growth and its potential inhibitory effects on cells were first evaluated. When 0.5–10 g/L 2-DG was supplied in the absence of glucose and xylose, no growth was observed (data not shown). This result could reflect either the inability of the strain to utilize 2-DG as a carbon source or growth inhibition caused by 2-DG itself. To distinguish these possibilities, the strain was cultivated in mP2 medium containing 15 g/L glucose and 0–10 g/L 2-DG. Growth profiles were similar across all tested 2-DG concentrations, reaching a maximum OD₆₀₀ of approximately 6.8 after 24 h (Fig. 1A). Thus, 2-DG did not appreciably inhibit growth when glucose was available. Together, these observations suggest that 2-DG did not support growth as the sole added sugar and that its presence did not cause a growth-inhibitory effect.

**Figure 1.**
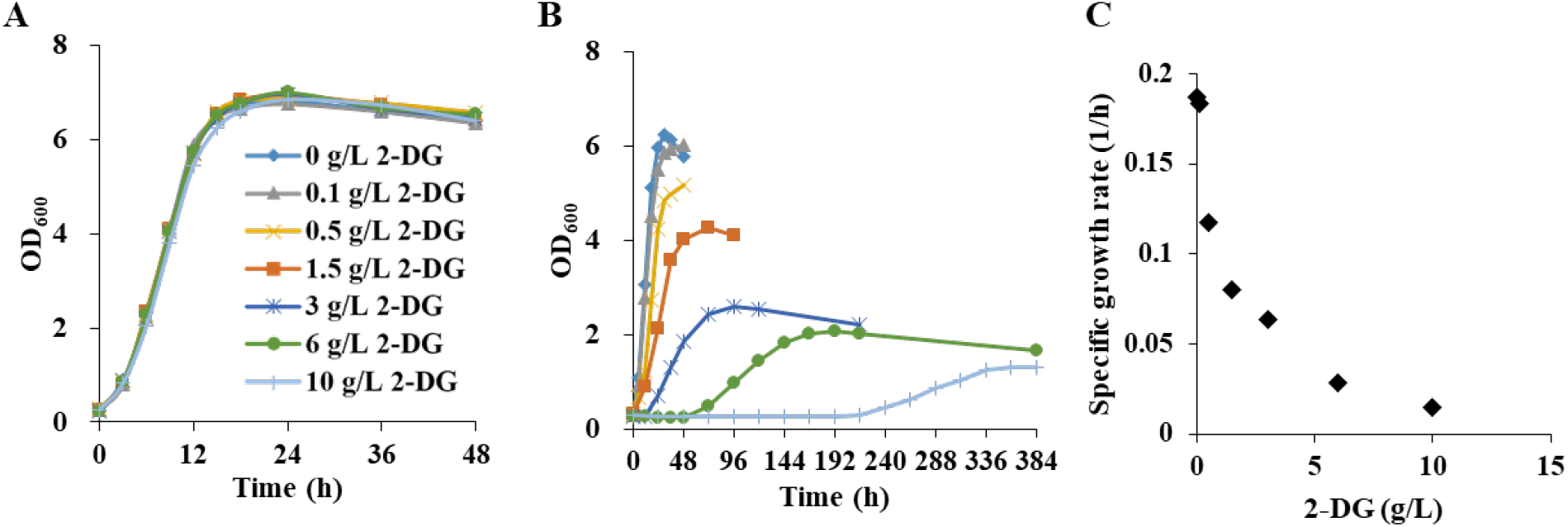
Effects of 2-deoxy-D-glucose (2-DG) on the growth of *C. tyrobutyricum* ATCC 25755 with glucose or xylose as the sole carbon source. Growth profiles in mP2 medium containing (A) 15 g/L glucose with 0–10 g/L 2-DG or (B) 15 g/L xylose with 0–10 g/L 2-DG. (C) Specific growth rates in mP2 medium containing 15 g/L xylose at different 2-DG concentrations. Cultures were incubated in serum bottles. Each fermentation was performed once (n = 1).

The effects of 2-DG were subsequently examined in mP2 medium containing 15 g/L xylose and 0–10 g/L 2-DG. Increasing 2-DG concentrations progressively delayed growth and reduced both the maximum OD₆₀₀ (Fig. 1B) and specific growth rate (Fig. 1C). In the absence of 2-DG, the specific growth rate was 0.187 h^-1^, and the maximum OD₆₀₀ of approximately 6.2 was reached after 30 h. At 0.5 and 1.5 g/L 2-DG, the specific growth rate decreased by approximately 37% and 57%, respectively. The maximum OD₆₀₀ decreased by approximately 17% and 31%, and the time required to reach these maxima increased to 48 and 72 h, respectively. A similar 2-DG concentration-dependent inhibition of growth on xylose was reported for the thermophilic Gram-positive bacterium *Parageobacillus thermoglucosidasius* DSM 2542 (Liang et al., 2022), in which increasing 2-DG concentrations reduced the peak OD₆₀₀ and delayed its occurrence in medium containing 10 g/L xylose.

The inhibition observed in xylose-containing medium, together with the absence of comparable inhibition in glucose-containing medium, suggests that 2-DG affected growth through interference with xylose utilization rather than through a general inhibitory effect on cells. This carbon source-dependent response is consistent with CCR-like inhibition of xylose utilization by a glucose analog.

### 3.2. Effect of 2-DG on growth and sugar consumption of *C. tyrobutyricum* in glucose-xylose mixtures

The inhibition of growth in xylose-containing medium suggested a CCR-like effect of 2-DG on xylose utilization (Section 3.1). To further characterize the CCR phenotype of *C. tyrobutyricum* ATCC 25755, growth and sugar consumption of the strain were examined in glucose–xylose mixtures with and without 2-DG. The 2-DG-containing culture was used to examine whether inhibition of xylose utilization persisted after glucose depletion while the glucose analog remained present. Low initial total sugar concentrations (1.3–2.1 g/L) were used to limit butyric acid accumulation and the accompanying pH decrease, thereby reducing their potential inhibitory effects on growth and sugar consumption.

In the absence of 2-DG (Fig. 2A), *C. tyrobutyricum* ATCC 25755 was capable of simultaneously utilizing 0.22 g/L of xylose while completely consuming 0.77 g/L glucose. After the depletion of glucose, a short growth lag of approximately 3 h was observed, after which growth continued and the remaining xylose was completely consumed (Fig. 2A). The partial co-utilization of glucose and xylose indicates that glucose did not completely suppress xylose utilization under these conditions. The consumption of xylose before glucose depletion agrees with previous observations of glucose–xylose co-utilization by *C. tyrobutyricum* (Oh et al., 2019; Wei et al., 2013; Zhu et al., 2002), supporting the interpretation that glucose-mediated inhibition was incomplete under the conditions tested.

**Figure 2.**
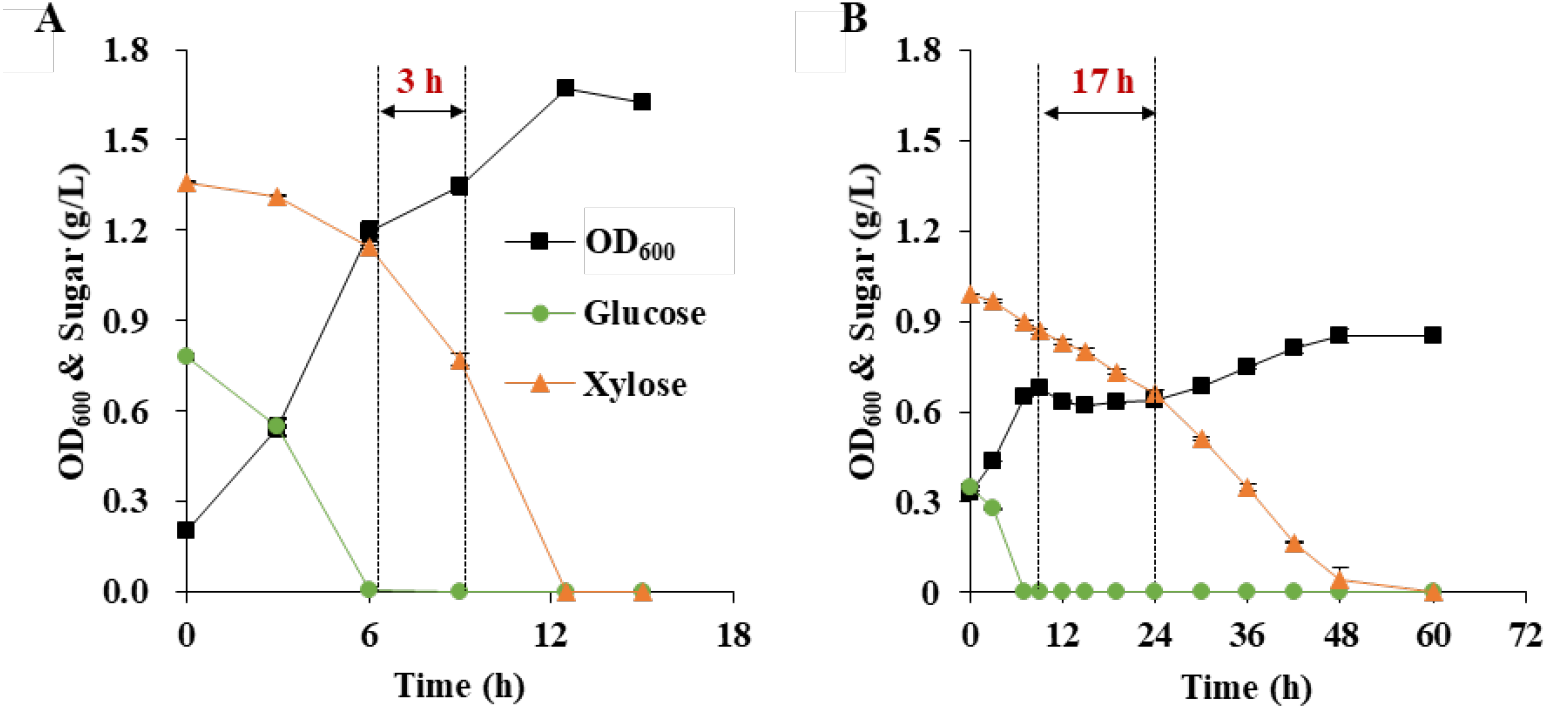
Effects of 2-deoxy-D-glucose (2-DG) on growth and sugar consumption of *C. tyrobutyricum* ATCC 25755 in glucose–xylose mixtures. Growth and sugar consumption profiles in mP2 medium containing glucose-xylose mixtures (A) without 2-DG and (B) with 0.5 g/L 2-DG. Double-headed arrows indicate the growth lag following glucose depletion. Data points and error bars represent the mean and range of duplicate experiments, respectively (n = 2).

In the presence of 0.5 g/L of 2-DG, *C. tyrobutyricum* ATCC 25755 consumed 0.10 g/L xylose while completely consuming 0.35 g/L glucose within 7 h (Fig. 2B). After glucose depletion, a diauxic-like growth lag of approximately 17 h was observed (Fig. 2B), compared with approximately 3 h in the absence of 2-DG (Fig. 2A). Nevertheless, xylose consumption continued during this period, and the remaining xylose was eventually depleted despite the continued presence of 2-DG. These results show that the 2-DG-containing culture exhibited a prolonged growth lag and slower xylose consumption, but xylose utilization was not completely blocked.

Together with the results in Section 3.1, these observations support a CCR-like inhibitory effect of 2-DG on xylose utilization. However, the consumption of xylose in the presence of either glucose or 2-DG indicates that the inhibition was incomplete under the conditions tested. These results suggest that xylose utilization by *C. tyrobutyricum* ATCC 25755 is not completely suppressed by glucose or 2-DG under the conditions tested.

### 3.3. Fermentation in different ratios of glucose and xylose by *C. tyrobutyricum* ATCC 25755

Glucose and xylose are major sugars obtained from lignocellulosic biomass, and efficient utilization of both sugars is important for maximizing biomass conversion (Olofsson et al., 2008). Glucose-to-xylose ratios in hydrolysates range from 3:1 to 1:9, depending on the feedstock and the pretreatment and hydrolysis conditions (Fu et al., 2022; Liang et al., 2013). Based on the CCR-like response observed in the preceding sections, the effects of glucose– xylose composition on sugar consumption and butyric acid production by *C. tyrobutyricum* ATCC 25755 were examined. Batch fermentations were conducted in a 2.5 L fermentor at pH 6.0 with a total initial sugar concentration of 80 g/L. Five sugar compositions were tested: 80 g/L glucose (G80), 80 g/L xylose (X80), 20 g/L glucose with 60 g/L xylose (G20X60), 40 g/L glucose with 40 g/L xylose (G40X40), and 60 g/L glucose with 20 g/L xylose (G60X20).

With either sugar supplied alone, glucose was consumed faster than xylose, with overall sugar consumption rates of 3.64 and 2.41 g/L/h, respectively (Fig. 3A, B; Table 1). The X80 culture also exhibited a longer initial lag phase than the G80 culture. The G80 culture produced 25.85 g/L butyric acid from 74.7 g/L glucose, with an overall productivity of 1.26 g/L/h. The X80 culture produced a higher final butyric acid titer of 28.64 g/L from 77.0 g/L xylose, but its overall productivity was lower, at 0.90 g/L/h, because of the longer fermentation time.

**Figure 3.**
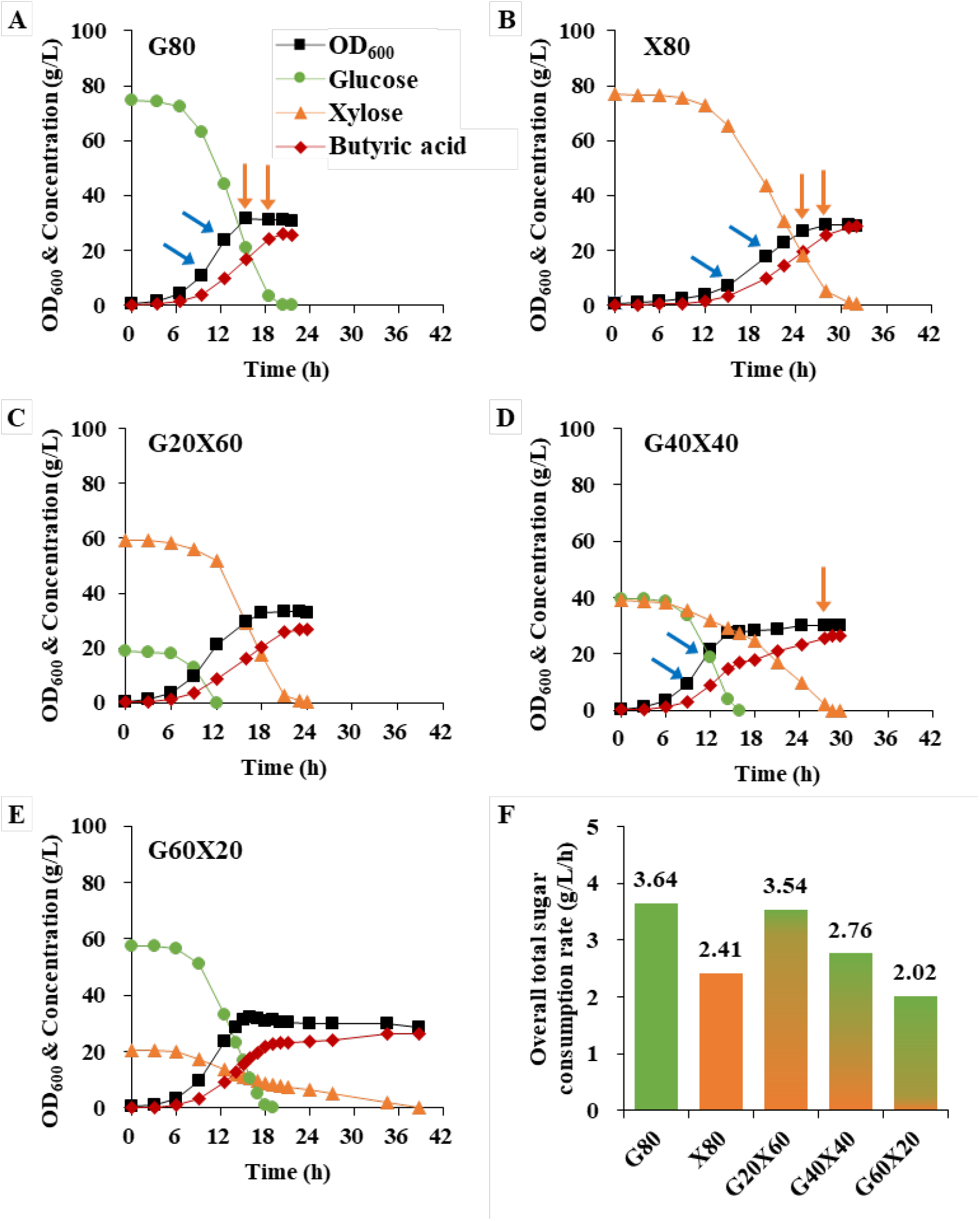
Effects of glucose–xylose composition on growth, sugar consumption, and butyric acid production by *C. tyrobutyricum* ATCC 25755. Growth, sugar consumption, and butyric acid production profiles in cultures supplied with (A) 80 g/L glucose (G80), (B) 80 g/L xylose (X80), (C) 20 g/L glucose and 60 g/L xylose (G20X60), (D) 40 g/L glucose and 40 g/L xylose (G40X40), and (E) 60 g/L glucose and 20 g/L xylose (G60X20). (F) Overall total sugar consumption rates under the different sugar compositions. Blue and orange arrows indicate sampling points for XylA activity measurements during the exponential and stationary phases, respectively. Each fermentation was performed once in a 2.5 L bioreactor (n = 1).

**Table 1.**
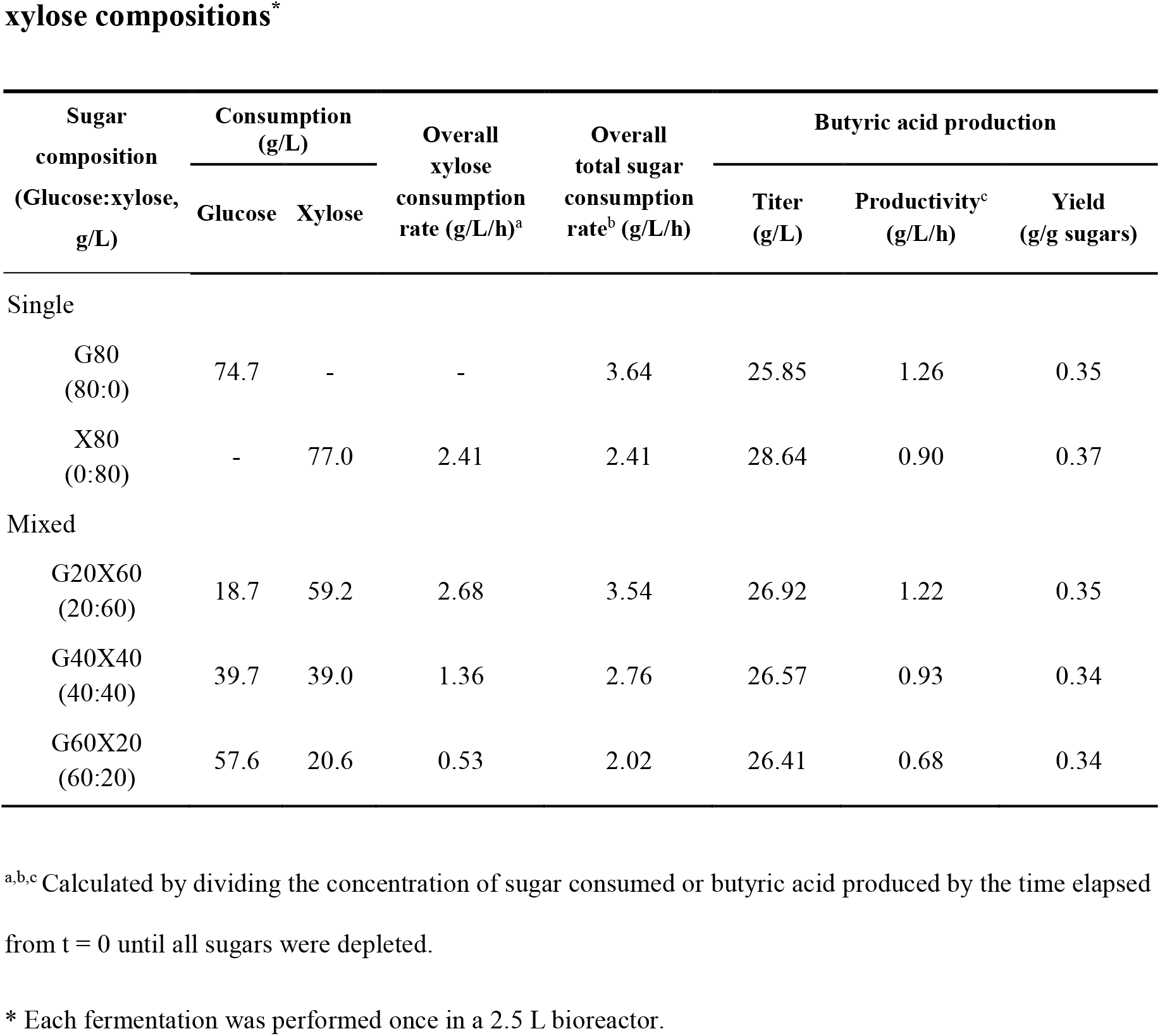
Fermentation performance of *C. tyrobutyricum* ATCC 25755 at different glucose-xylose compositions*.

In all mixed-sugar cultures, glucose was consumed preferentially during the initial growth period, accompanied by consumption of a small amount of xylose (Fig. 3C–E). The overall xylose consumption rates decreased as the initial glucose proportion increased, from 2.68 g/L/h in G20X60 to 1.36 g/L/h in G40X40 and 0.53 g/L/h in G60X20. Consequently, complete xylose consumption took the longest in G60X20 despite its lowest initial xylose concentration. These patterns suggest stronger suppression of xylose utilization in mixtures containing a higher proportion of glucose.

Interestingly, the overall xylose consumption rate in G20X60 was slightly higher than that in X80, despite the CCR-like phenotype observed in the preceding experiments. Moreover, despite its high xylose proportion, G20X60 exhibited an overall total sugar consumption rate of 3.54 g/L/h, close to the 3.64 g/L/h obtained in G80 (Fig. 3F; Table 1). This performance may reflect a combination of less pronounced glucose-mediated inhibition at the relatively low initial glucose concentration and early biomass formation supported by glucose. The latter could help offset the longer initial lag observed in X80, although the contribution of each factor remains to be established.

Among the mixed-sugar cultures, G20X60 achieved the highest butyric acid productivity, at 1.22 g/L/h (Table 1). Notably, despite its high xylose proportion, its productivity was close to that obtained with glucose alone in G80 (1.26 g/L/h) and was approximately 1.36 times that in X80 (0.90 g/L/h). Butyric acid titers and yields were similar across the mixed-sugar conditions, ranging from 26.41 to 26.92 g/L and from 0.34 to 0.35 g/g sugars, respectively (Table 1). Thus, sugar composition primarily affected fermentation time and productivity rather than final titer or yield. These results suggest that a xylose-rich mixture containing a relatively low proportion of glucose may support fermentation performance comparable to that obtained with glucose alone, despite preferential glucose consumption.

Fu et al. (2022) proposed that HPrK-dependent phosphorylation of HPr enables the formation of a CcpA–HPr(Ser-P) complex, which represses the transcription of *xylAB* in *C. tyrobutyricum*. Because a small amount of xylose was consumed while glucose remained present in the mixed-sugar cultures in the present study (Fig. 3), the activity of XylA (xylose isomerase encoded by *xylA*), which catalyzes the conversion of xylose to xylulose, was examined to determine whether it remained detectable in the presence of glucose. In G40X40, XylA activity was measured (i) during the exponential phase, when glucose and xylose were simultaneously consumed, and (ii) during the stationary phase, after glucose depletion, when xylose remained available. For comparison, XylA activity in G80 and X80 cultures was also measured during the exponential and stationary phases.

In the single-sugar cultures, XylA specific activity was similar between the exponential and stationary phases (Fig. 4). Activity was substantially higher in cells grown on xylose (10.74 U/mg protein) than in those grown on glucose (1.05 U/mg protein). In G40X40, XylA activity was 4.52 U/mg protein during the exponential phase and increased to 7.85 U/mg protein after glucose depletion, representing a 1.74-fold increase. The detectable activity while glucose was present, together with its increase after glucose depletion, is consistent with partial suppression of xylose isomerase activity during glucose–xylose co-utilization. These enzyme activity results support the sugar consumption patterns in Fig. 3 and suggest that glucose-mediated inhibition of xylose metabolism was incomplete under the conditions tested.

**Figure 4.**
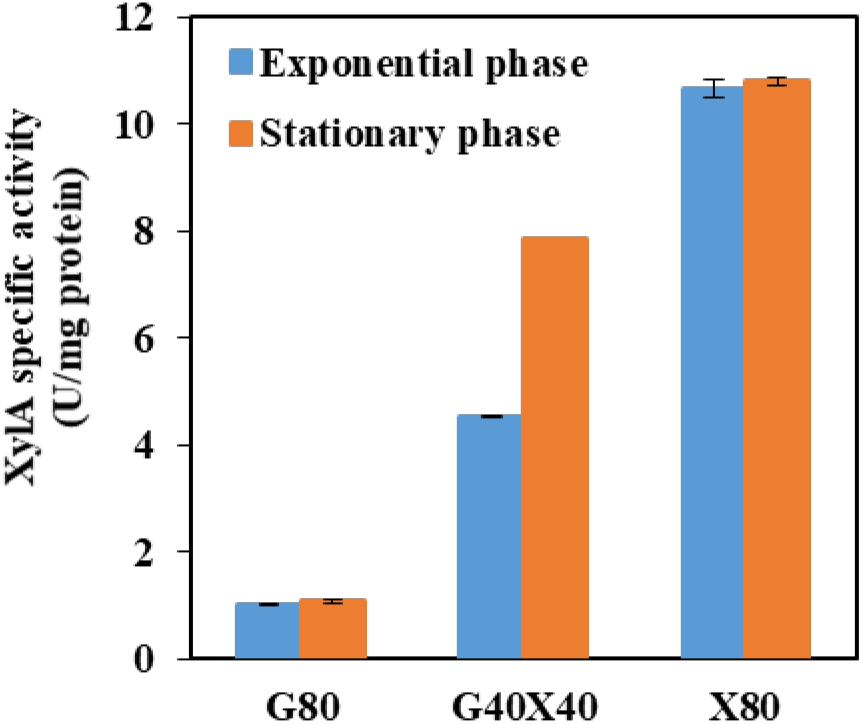
Xylose isomerase (XylA) specific activity in *C. tyrobutyricum* ATCC 25755 grown on glucose, xylose, or a glucose-xylose mixture. Specific XylA activity was measured in cells collected during the exponential and stationary phases of cultures supplied with 80 g/L glucose (G80), 40 g/L glucose and 40 g/L xylose (G40X40), or 80 g/L xylose (X80). Sampling points are indicated in Fig. 3A, 3B, and 3D. One unit of activity was defined as the production of 1 µmol xylulose per minute, and specific activity was expressed as U/mg protein. Bars and error bars represent the mean and range, respectively, of XylA activities measured in samples collected at two different time points during each growth phase, except for the stationary phase of G40X40. For this condition, the bar represents activity measured in a sample collected at a single time point, and no error bar is shown.

### 3.4. Fermentation of *C. tyrobutyricum* ATCC 25755 on a rice straw hydrolysate

The G20X60 culture showed rapid xylose consumption after glucose depletion and fermentation performance comparable to that obtained with glucose alone (Fig. 3C; Table 1). To examine whether a similar sugar consumption pattern could be observed with a lignocellulosic substrate, *C. tyrobutyricum* ATCC 25755 was cultivated in mP2 medium supplemented with rice straw hydrolysate prepared by dilute-acid hydrolysis. The initial glucose and xylose concentrations were 5.0 and 15.1 g/L, respectively, corresponding to a glucose-to-xylose ratio of approximately 1:3, similar to that in G20X60.

Both sugars were completely consumed within approximately 16 h, producing 7.68 g/L butyric acid with an overall productivity of 0.48 g/L/h and a yield of 0.38 g/g sugars (Fig. 5). The overall total sugar consumption rate was 1.26 g/L/h. Although the glucose-to-xylose ratio was similar to that in G20X60, the overall sugar consumption rate and butyric acid productivity were lower in the hydrolysate culture. This difference may partly reflect the effects of hydrolysate-associated inhibitors. Compounds such as furfural, 5-hydroxymethylfurfural, and phenolics can be generated during acid pretreatment and hydrolysis. Previous studies have reported that hydrolysate-derived phenolic compounds inhibited *C. tyrobutyricum* fermentation, resulting in a prolonged lag phase and slower sugar consumption (Lee et al., 2015; Luo et al., 2024). However, the contribution of these compounds to the present results remains uncertain, and the difference in total initial sugar concentration between the hydrolysate culture and G20X60 should also be considered.

**Figure 5.**
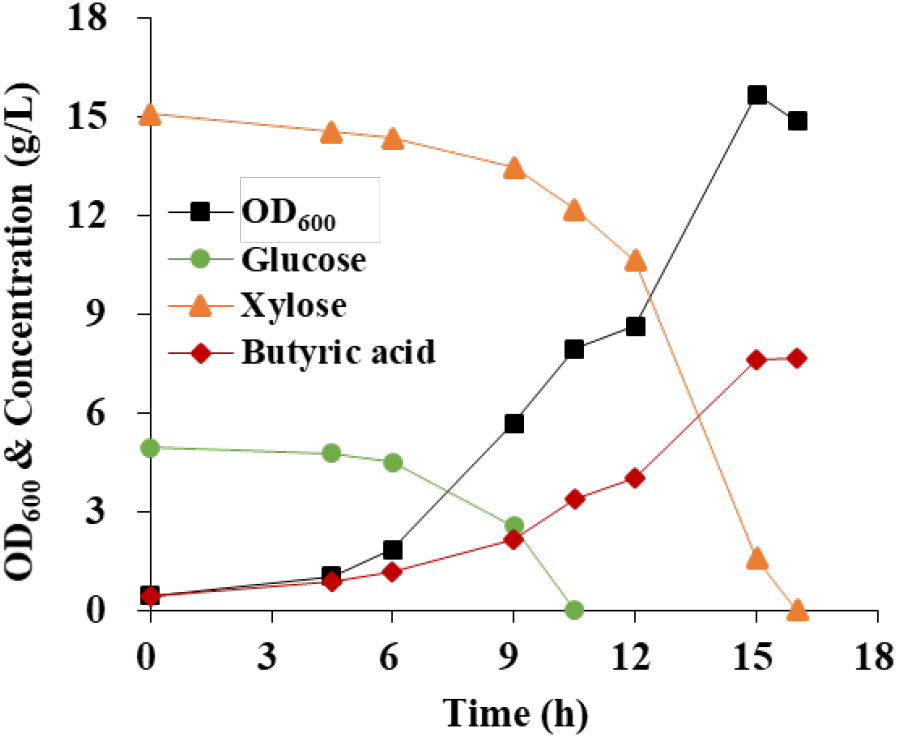
Growth, sugar consumption, and butyric acid production by *Clostridium tyrobutyricum* ATCC 25755 during fermentation of xylose-rich rice straw hydrolysate. Fermentation was performed once in a 2.5 L bioreactor (n = 1).

Despite the lower overall fermentation rates, xylose was rapidly consumed after glucose depletion, following a pattern similar to that observed in G20X60 (Fig. 3C). This result suggests that xylose utilization was not subject to sustained inhibition in the xylose-rich hydrolysate under the conditions tested. Together with the results obtained using synthetic sugar mixtures, this observation supports the applicability of *C. tyrobutyricum* ATCC 25755 to substrates containing a relatively low proportion of glucose and a high proportion of xylose.

Depending on the pretreatment conditions, lignocellulosic biomass can be fractionated into a hemicellulose-derived, xylose-rich liquid fraction and a cellulose-rich solid fraction. The latter can subsequently be enzymatically hydrolyzed to obtain glucose for ethanol production and other fermentation processes. Utilization of the xylose-rich liquid fraction may be more challenging because of inhibitory compounds and the limited xylose utilization capacity of some microorganisms. Nevertheless, xylose-rich hydrolysates derived from bioenergy sorghum, spent lemongrass, and rice straw have been used to produce β-carotene, clavulanic acid, and biosurfactants, respectively (Cheng et al., 2020; Kumar et al., 2023; Panjiar et al., 2020).

The results of the present study suggest that *C. tyrobutyricum* ATCC 25755 can effectively utilize glucose–xylose mixtures under xylose-rich conditions without genetic modification to eliminate CCR. This capability is supported by its effective xylose utilization (Fig. 3C) and the limited glucose-mediated inhibition observed under the xylose-rich conditions tested. These findings suggest the potential of this strain for valorizing xylose-rich lignocellulosic fractions through butyric acid production. Further optimization of hydrolysate preparation and fermentation conditions may help improve sugar consumption rates and butyric acid productivity.

## 4. Conclusions

In mixed-sugar fermentations, increasing the glucose proportion was associated with slower xylose consumption by *C. tyrobutyricum* ATCC 25755. However, partial glucose– xylose co-utilization and detectable XylA activity suggested that glucose did not completely repress xylose metabolism. The xylose-rich G20X60 culture achieved an overall total sugar consumption rate of 3.54 g/L/h and a butyric acid productivity of 1.22 g/L/h, comparable to those obtained with glucose alone. These results suggest that a high xylose proportion can support fermentation performance comparable to that achieved with glucose alone, despite the CCR-like behavior observed in this strain. This potential was further supported by fermentation of xylose-rich rice straw hydrolysate with a glucose-to-xylose ratio of 1:3, similar to that in G20X60. Together, these findings support the potential of wild-type *C. tyrobutyricum* ATCC 25755 for utilizing xylose-rich lignocellulosic hydrolysates for butyric acid production.

## CRediT author statement

**Ki-Yeon Kim**: Writing–original draft, Methodology, Visualization, Investigation. **Hyun Ju Oh**: Writing–original draft, Data curation, Methodology. **Jung Ho Ahn**: Writing – review & editing, Supervision. **Ja Kyong Ko**: Resources, Supervision. **Gyeongtaek Gong**: Writing – review & editing, Validation. **Youngsoon Um**: Writing – review & editing, Supervision, Project administration, Funding acquisition.

## Data availability

Data will be made available on request.

